# T cell responses towards PINK1 and α-synuclein are elevated in prodromal Parkinson’s disease

**DOI:** 10.1101/2025.04.21.649871

**Authors:** Emil Johansson, Antoine Freuchet, Gregory P. Williams, Tanner Michealis, April Frazier, Irene Litvan, Jennifer G Goldman, Roy N Alcalay, David G. Standaert, Amy W. Amara, Natividad Stover, Edward A. Fon, Ronald B. Postuma, John Sidney, David Sulzer, Cecilia S. Lindestam Arlehamn, Alessandro Sette

## Abstract

A role of the immune system in Parkinson’s disease (PD) progression has long been suspected due to the increased frequency of activated glial cells and infiltrating T cells into the substantia nigra. It was previously reported that PD donors have increased T cell responses towards PINK1 and α-synuclein (α-syn), two Lewy body-associated proteins. Further, T cell reactivity towards α-syn was highest closer to disease onset, highlighting that autoreactive T cells might play a role in PD pathogenesis. However, whether T cell autoreactivity is present during prodromal PD is unknown. Here, we investigated T cell responses towards PINK1 and α-syn in donors at high risk of developing PD (i.e. prodromal PD: genetic risk, hyposmia, and or REM sleep behavior disorder), in comparison to PD and healthy control donors. T cell reactivity to these two autoantigens was detected in prodromal PD at levels comparable to those detected in individuals with clinically diagnosed PD. Aligned with the increased incidence of PD in males, we found that males with PD, but not females, had elevated T cell reactivity compared to healthy controls. However, among prodromal PD donors, males and females had elevated T cell responses. These differing trends in reactivity highlights the need for further studies of the impact of biological sex on neuroinflammation and PD progression.

## Main

Parkinson’s Disease (PD) is associated with accumulation of Lewy bodies (LBs), primarily composed of α-synuclein (α-syn), as well as other proteins such as PINK^1^. LBs disrupt cellular processes in neurons, and eventually cause a loss of neurons ^2^. In addition to cytotoxic effects on neurons^3^, LBs can also induce neuroinflammation by activating glial cells^4^. Several lines of evidence support a role for neuroinflammation and autoimmunity in PD, such as observations of increased frequencies of activated glial cells^5^ and infiltrating T cells in the substantia nigra pars compacta (SNpc)^6, 7^. We previously reported that PD patients have increased frequencies of T cells recognizing PINK1 and α-syn compared to healthy controls (HC)^8-10^, suggesting that autoimmunity is a feature of PD. For reasons still unknown, males are at a higher risk of developing PD^11^. In line with this, we previously found that males and females with PD and HCs can have different patterns of reactivity towards different neuroantigens^9^. However, it is still unknown which role these differences in immune responses play in PD pathology, and if these differences are induced prior to PD diagnosis and contribute to PD pathogenesis.

Although motor symptoms are the hallmark symptoms of PD, non-motor symptoms such as hyposmia/anosmia, REM sleep behavior disorders (RBD), and gastrointestinal dysfunction often precede the PD diagnosis^12^. It is widely recognized that PD pathogenesis is initiated long before diagnosis, in a prodromal phase that may take decades^12^. We previously reported a case study of a participant for which stored blood samples were available for study, collected almost a decade before and a decade after diagnosis. T cell reactivity towards α-syn was highest closest to motor disease onset^8^. Whether autoimmune T cell reactivity is present during the prodromal phase is an important open question. Understanding early drivers of PD disease progression could identify prognostic markers to identify individuals at risk of developing PD and guide the development of therapeutic interventions.

Here we investigated if T cell responses towards PINK1 and α-syn are present during the prodromal phase of PD in a group of donors at a high risk of developing PD, and assessed the impact of sex on T cell responses before and after PD onset. The study included 82 prodromal donors, including individuals with gene mutations linked to increased risk of developing PD (*GBA* and *LRRK2;* n=27), individuals diagnosed with hyposmia (n=29), a combination of hyposmia and mutations (*SNCA, GBA*, or *LRRK2*, n=3), individuals diagnosed with RBD (n=19), or a combination of RBD and hyposmia (n=4). The study also included 70 age- and sex-matched HC and 70 age-matched individuals with PD (**Table 1**). The PD group had significantly fewer female donors, reflecting the typical lower PD incidence of females. All three groups were of similar age, but PD donors have significantly higher Unified Parkinson’s Disease Rating Scale, part III scores, reflecting their motor function impairments. Importantly, no difference in age or motor function was identified between male and female donors within each group.

**Table 1.**
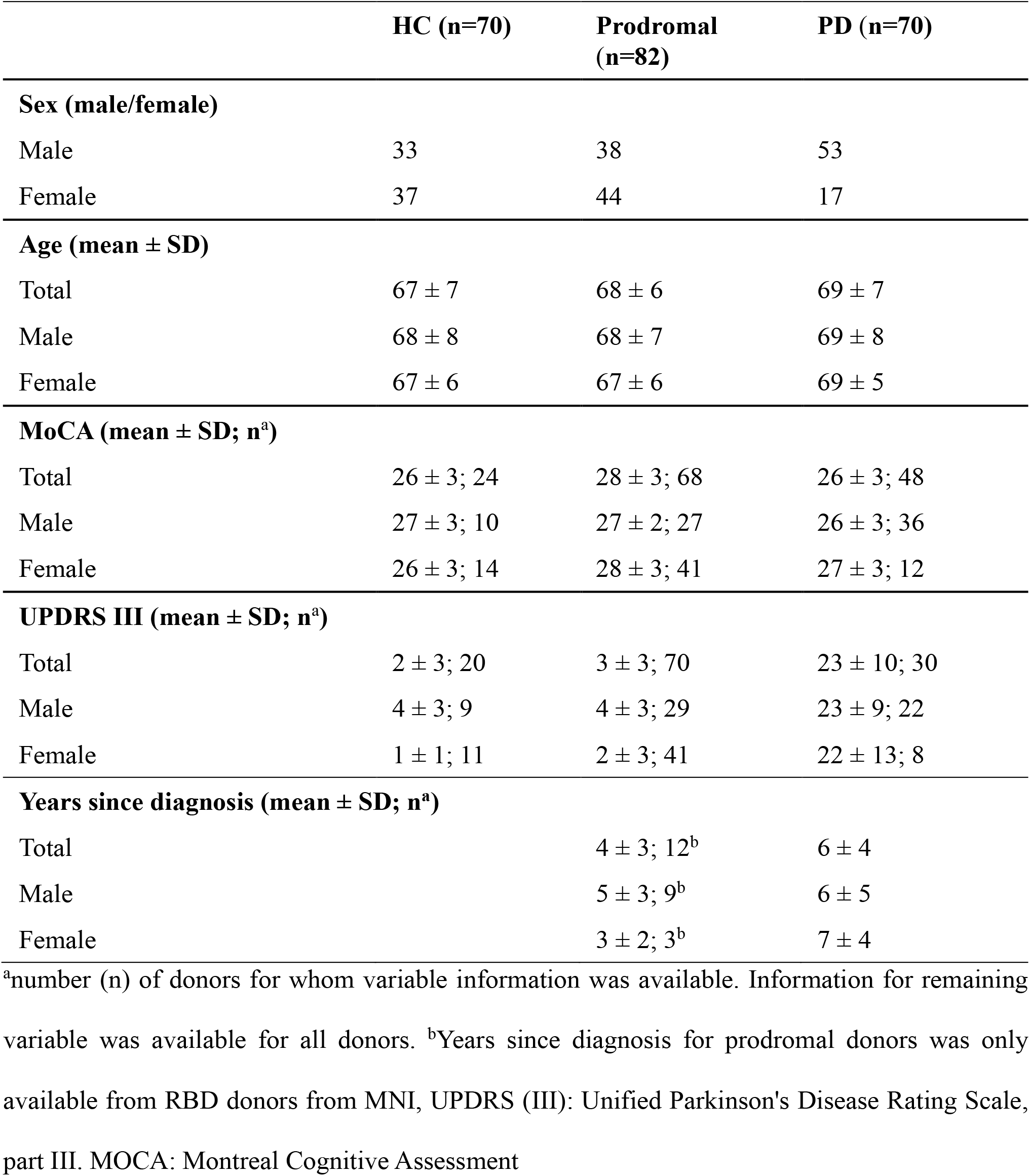
Characteristics of the study cohorts.

|  | **HC (n=70)** | **Prodromal**<br>**(n=82)** | **PD (n=70)** |
| --- | --- | --- | --- |
| **Sex (male/female)** |  |  |  |
| Male | 33 | 38 | 53 |
| Female | 37 | 44 | 17 |
| **Age (mean ± SD)** |  |  |  |
| Total | 67 ± 7 | 68 ± 6 | 69 ± 7 |
| Male | 68 ± 8 | 68 ± 7 | 69 ± 8 |
| Female | 67 ± 6 | 67 ± 6 | 69 ± 5 |
| **MoCA (mean ± SD; n<sup>a</sup>)** |  |  |  |
| Total | 26 ± 3; 24 | 28 ± 3; 68 | 26 ± 3; 48 |
| Male | 27 ± 3; 10 | 27 ± 2; 27 | 26 ± 3; 36 |
| Female | 26 ± 3; 14 | 28 ± 3; 41 | 27 ± 3; 12 |
| **UPDRS III (mean ± SD; n<sup>a</sup>)** |  |  |  |
| Total | 2 ± 3; 20 | 3 ± 3; 70 | 23 ± 10; 30 |
| Male | 4 ± 3; 9 | 4 ± 3; 29 | 23 ± 9; 22 |
| Female | 1 ± 1; 11 | 2 ± 3; 41 | 22 ± 13; 8 |
| **Years since diagnosis (mean ± SD; n<sup>a</sup>)** |  |  |  |
| Total | | 4 ± 3; 12<sup>b</sup> | 6 ± 4 |
| Male | | 5 ± 3; 9<sup>b</sup> | 6 ± 5 |
| Female | | 3 ± 2; 3<sup>b</sup> | 7 ± 4 |

<sup>a</sup>number (n) of donors for whom variable information was available. Information for remaining variable was available for all donors. <sup>b</sup>Years since diagnosis for prodromal donors was only available from RBD donors from MNI, UPDRS (III): Unified Parkinson's Disease Rating Scale, part III. MOCA: Montreal Cognitive Assessment

To identify α-syn and PINK1 reactive T cells we first stimulated PBMCs with previously described peptide pools^9, 10^ to expand antigen-specific T cells for two weeks^9, 10^, and a Fluorospot assay was used to quantify the number of IFNγ- or IL-5-producing T cells following restimulation with the peptide pools. A subset of the HC and PD donors were previously screened for reactivity towards PINK1^9^, and data from the individuals screened for both α-syn and PINK1 is included here for comparison, while the prodromal cohort data is novel. We also included an Epstein-Barr virus (EBV) peptide pool^13^ as a specificity control. As in the original paper^9^, PD donors had significantly higher IFNγ and IL-5 responses towards PINK1 compared to HCs (p=0.02, and p=0.03, respectively; **Fig 1A**). Further, the results confirm our previous findings that PD donors had significantly higher IFNγ and IL-5 responses towards α-syn compared to HCs (p=0.01, and p=0.02, respectively; **Fig 1B**), but not to EBV (**Supplemental Figure S1**). The PINK1 and α-syn reactivity of prodromal donors was at levels similar to the individuals with PD for both IFNγ and IL-5 mediated T cell responses. Intriguingly, IFNγ and IL-5 reactivity towards PINK1 was significantly higher in prodromal than in HC donors (p=0.02, and p=0.003, respectively; **Fig 1A**). Similarly, IL-5 responses to α-syn were increased in prodromal donors (p=0.05; **Fig 1B**), and a trend was also observed for increased IFNγ responses to α-syn in prodromal donors (p=0.1; **Fig 1B**), compared to HCs. The current sample size did not allow for a properly statistically powered comparison of the T cell responses towards PINK1 and α-syn between the different prodromal subgroups (**Supplemental Fig 1**), or T cell reactivity and time since diagnosis in RBD and PD donors (**Supplemental Fig 3A-C**).

**Figure 1.**
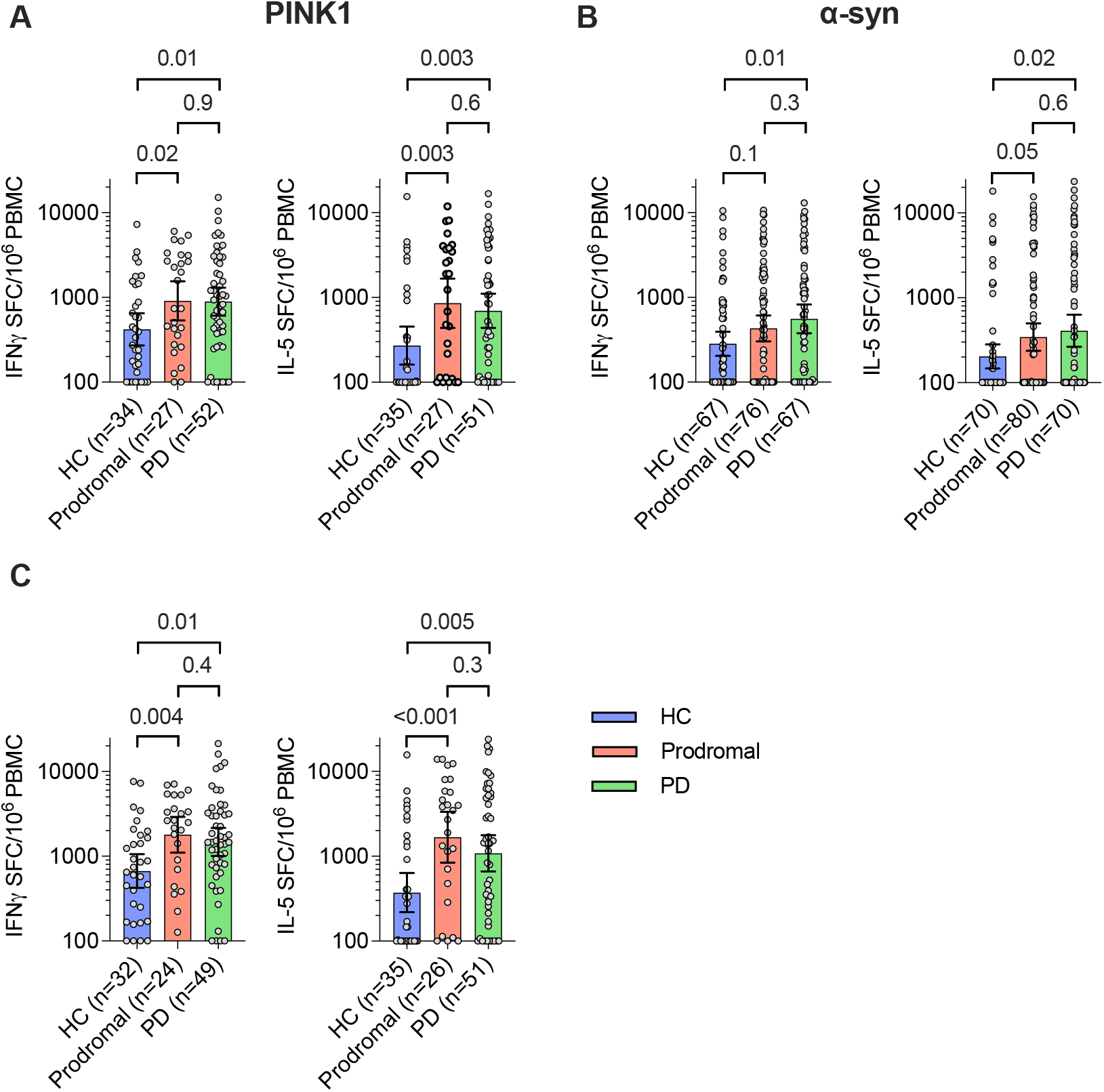
PD and prodromal donors have elevated T cell responses towards PINK1 and α- syn. Magnitude of IFNγ and IL-5 mediated T cell responses towards PINK1 (**A**) and α-synuclein (α-syn; **B**) in healthy controls (HC), prodromal, and Parkinson’s disease (PD) donors. Magnitude of pooled IFNγ (left panel) and IL-5 (right panel) responses to PINK1 and α-syn (**C**), in donors screened for reactivity towards both peptide pools. Each circle represents an individual donor. P-values from two-tailed Kruskal-Wallis tests followed by an uncorrected Dunns’ test, and geometric mean ± 95% confidence interval are shown.

We note that not all PD donors respond to α-syn^14^, and it has been hypothesized that this might reflect the kinetics of responses associated with disease progression, as well as the presence of additional antigens associated with PD pathogenesis. Indeed, the combined IFNγ- and IL-5- mediated T cell responses towards PINK1 and α-syn were higher in both prodromal (p=0.004, p<0.001, respectively) and PD (p=0.01, p=0.005, respectively) donors than HC donors (**Fig 1C**). Taken together, these results demonstrate that prodromal PD donors have increased T cell responses to two previously described targets of autoreactive T cells in PD, indicating that autoimmune responses are present during the prodromal PD disease stage.

We previously reported biological sex differences in autoantigen reactivity in PD^9^. While the statistical power of the analysis is limited by the number of observations in each subgroup, we found that male PD donors had higher IFNγ-mediated T cell responses to PINK1 and α-syn (p=0.01, p=0.03, respectively; **Fig 2A, B**), higher IL-5 responses to PINK1 and a trend of higher α-syn (p=0.007 and p=0.07; **Fig 2C, D**), compared to male HCs. In contrast, female PD donors did not have elevated T cell responses towards PINK1 or α-syn compared to female HCs, and had significantly lower IFNγ-mediated T cell responses to α-syn compared to male PD (p=0.019; **Supplemental Figure S3D, E**). Among prodromal donors, both male and female subjects had higher IL-5 responses to PINK1 than HC (p=0.04, p=0.03, respectively; **Fig 2C**), and female prodromal donors had higher IFNγ responses to PINK1 than HC donors (p=0.02; **Fig 2A**).

**Figure 2.**
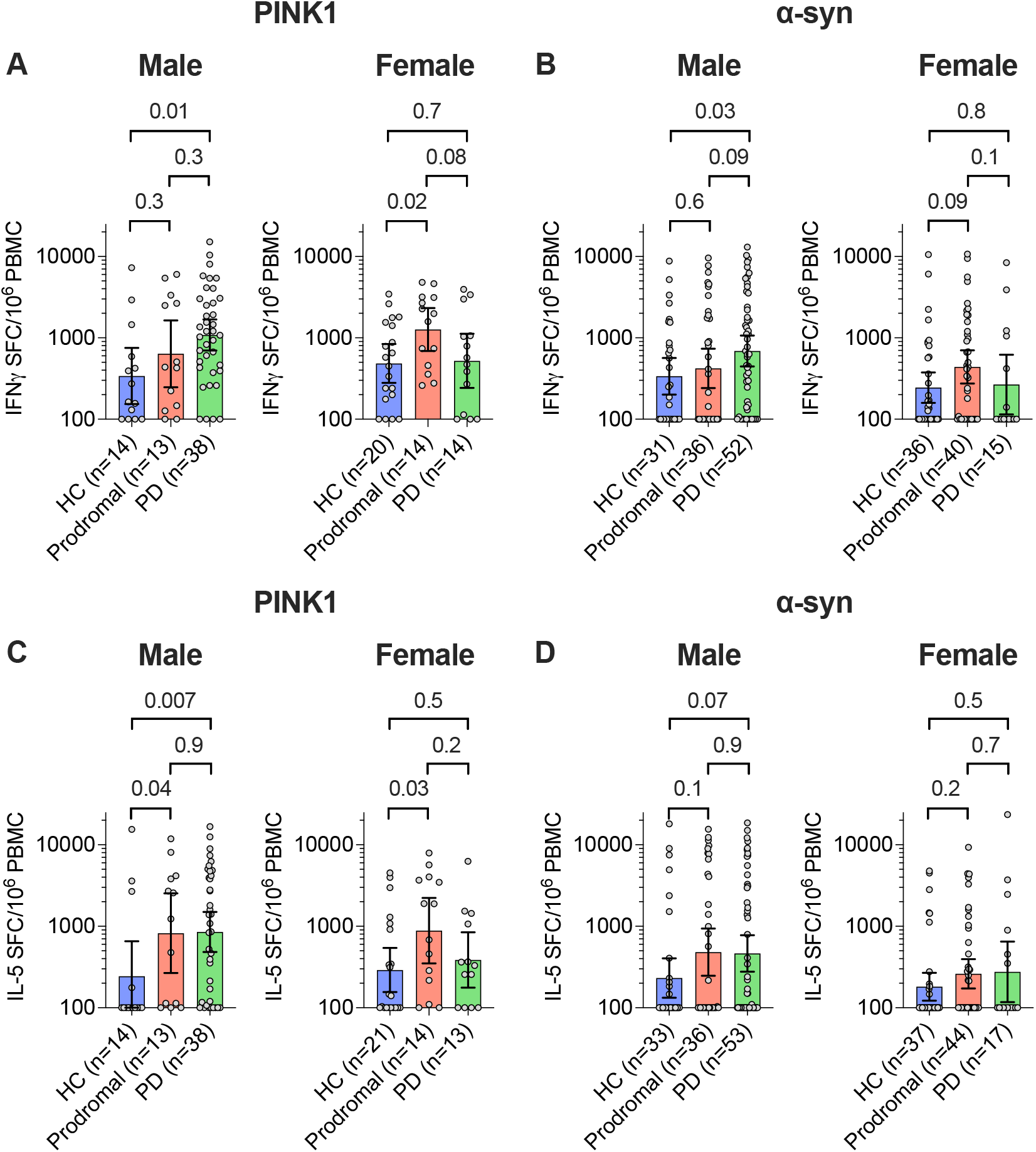
T cell reactivity towards PINK1 is increased in both male and female prodromal donors. Magnitude of IFNγ responses towards PINK1 (**A**) and α-synuclein (α-syn; **B**), and IL-5 responses towards PINK1 (**C**) and α-syn (**D**), in male (left panels) and female (right panel) separated. Each circle represents an individual donor. P-values from two-tailed Kruskal-Wallis tests followed by an uncorrected Dunns’ test, and geometric mean ± 95% confidence interval are shown.

In conclusion, this study confirms that α-syn T cell reactivity is associated with PD. In addition, we found that PINK1 and α-syn reactivity is also detectable in the prodromal phase at levels comparable to that seen in PD cases, in line with previous reports of increased neuroinflammation and microglial activation in prodromal PD donors^15^. It will be of interest for future studies to determine if the increased frequencies of activated T cells and monocytes expressing transmigration receptor reported in RBD donors^15^ are associated with autoimmune T cell responses. The pathogenic role of CD4 T cells in PD is believed to, in part, be mediated by direct toxic effect on neurons^16^, and by contributing to microgliosis^17-19^. We further report biological sex differences in prodromal IFNγ and IL-5 reactivity towards PINK1 in males and females with PD. The molecular mechanism for the increase in PINK1 reactivity in both male and female prodromal donors, but only in male PD donors, noted here is unclear, and the influence of biological sex on immune perturbations during disease progression in PD, and other neurodegenerative diseases, is still poorly known^20^. The observation that prodromal female, but not female PD, subjects have increased frequency of autoreactive T, is in contrast with the lower disease incidence among women, and raises the intriguing possibility that females in prodromal stages develop similar levels of α-syn T cell reactivity to PD males, but do not progress to diagnosed PD. This is in contrast to relapsing remitting multiple sclerosis (RRMS), where strong autoimmune T cell responses are believed to contribute to the increased incidence among females^21^. This could be related to the observation that healthy female donors have lower *PINK1* and *SNCA* gene expression in the SNpc^22^, thereby limiting the detrimental effect of these cells in female prodromal donors, or that diagnosis in females occurs later than in males^11^ after the overall number of T cells is decreased.

Although these donors are at high risk of developing PD, we do not know yet if and when these donors will develop PD. In addition to PD, these donors are also at risk of developing other synucleinopathies, such as dementia with Lewy bodies (DLB) and multiple system atrophy (MSA)^23^. It will therefore be of great interest to further study these donors longitudinally to assess the relationship between neuroantigen-specific T cell reactivity and time to PD, DLB, or MSA diagnosis. The observation of neuroantigen-specific T cell reactivity in the prodromal phase is consistent with the intriguing observation that individuals who are affected inflammatory bowel diseases and are treated with anti-inflammatory agents such as anti-TNF antibodies have a lower incidence of PD^24^. In conclusion, our results support the hypothesis that detecting early T cell responses might aid in early diagnosis and that interfering with inflammation in the prodromal phase might positively impact PD disease progression.

## Materials and methods

### Study approval

All participants provided written informed consent for participation in the study. Ethical approval was obtained from the Institutional Review Boards at LJI (protocol numbers: VD-124 and VD-118), CUMC (protocol number IRB-AAAQ9714 and AAAS1669), UCSD (protocol number 161224), Shirley Ryan Ability Lab/Northwestern University (protocol number STU00209668-MOD0005), the Parkinson’s Progression Markers Initiative (PPMI; protocol number 20216216 and 20200597), University of Alabama at Birmingham (UAB; protocol number IRB-300001297), and Montreal Neurological Institute (MNI; protocol number 2017-330, 15-944-MUHC).

### Study participants

Subjects with PD (n=45), prodromal donors with RBD (n=12), and HCs (n=59) were recruited by the Movement Disorders Clinic at the Department of Neurology at CUMC, by the clinical core at LJI, by the Parkinson and Other Movement Disorder Center at UCSD, by the movement disorder specialists at the Parkinson’s disease and Movement Disorders program at Shirley Ryan Ability Lab, by movement disorder specialists at UAB, and from the MNI. In addition, the PPMI (https://dx.doi.org/10.17504/protocols.io.n92ldmw6ol5b/v2) also provided samples collected from 11 HC, 70 prodromal, and 25 PD donors. For the 16 PD donors recruited from LJI, PD disease status and medical history was self-reported. Inclusion criteria for the remaining 54 PD patients consisted of (I) clinically diagnosed PD with the presence of bradykinesia and either resting tremor or rigidity, (II) PD diagnosis between ages 35–80, (III) history establishing dopaminergic medication benefit, and (IV) ability to provide informed consent. Exclusion criteria for PD were atypical parkinsonism or other neurological disorders, history of cancer within past 3 years, autoimmune disease, and chronic immune modulatory therapy. Age-matched HCs were selected on the basis of (I) age 45–85 and (II) ability to provide informed consent. Exclusion criteria for HCs were the same as PD except for the addition of self-reported PD genetic risk factors (i.e., PD in first-degree blood relative). Tests for hyposmia was not performed for the HC cohort, and we are therefore unable to exclude the possibility of included HC donors with undiagnosed hyposmia^25^. The recruited individuals with PD all met the UK Parkinson’s Disease Society Brain Bank criteria for PD. Cohort characteristics are shown in Table 1. For a subset of participants, Movement Disorder Society–Unified Parkinson’s Disease Rating Scale, part III (MDS-UPDRS (III)) and Montreal Cognitive Assessment (MoCA) information was collected. The MDS-UPDRS (III) is a standard scale used to assess the motor function of PD patients, with higher numbers reflecting a greater loss of motor function. The MoCA score is a standard test for cognitive assessment, with a score of 26–30 points representing normal cognition and lower scores representing cognitive impairment.

### Sex as a biological variable

Our study included both male and female participants. The results have been reported as an aggregate for the entire cohort, and additionally, with female and male participants analyzed separately.

### PBMC isolation

Whole blood samples were collected in either heparin or EDTA containing blood bags or tubes. PBMCs were subsequently isolated using Ficoll-Paque Plus (Cytavia) according to the manufacturer’s instruction, as detailed in our published protocol (https://dx.doi.org/10.17504/protocols.io.bw2ipgce). Briefly, whole blood was first centrifuged to allow the collection of plasma, and the serum-depleted blood was thereafter diluted in RPMI, layered on Ficoll-Paque Plus, and centrifuged at 803g for 25 minutes with the brakes off. The interphase cell layer resulting from this spin was collected, washed with RPMI, counted, and cryopreserved in 90% v/v FBS and 10% DMSO and stored in liquid nitrogen until tested.

### Peptide pools

Previously defined 15-mer peptides derived from PINK1^9^, α-syn^8^, and EBV^13^ were used in this study. Peptides were synthesized commercially as crude material on a 1 mg scale by TC Peptide Lab. Lyophilized peptide products were dissolved in 100% (DMSO) at a concentration of 20 mg/mL, and their quality was spot-checked by mass spectrometry. The purity is greater than 85% for more than 85% of the peptides. Peptides were combined into megapools^26^, and used for antigen stimulation experiments.

### *In vitro* expansion and quantification of antigen specific cells

Antigen-specific cells were expanded and quantified as previously described^8, 9^ and detailed in our published protocol (https://www.protocols.io/view/pbmc-stimulation-with-peptide-pools-and-fluorospot-bphjmj4n). Briefly, PBMCs were thawed and then stimulated with megapools (5 μg/mL) for 4 days. After 4 days, cells were supplemented with fresh RPMI and IL-2 (10 U/mL, ProSpec Bio) and thereafter every 3 days, cells were supplied with fresh RPMI and IL-2. After 14 days, IFN-γ and IL-5 T cell responses to megapools were measured using a FluoroSpot assay (Mabtech FSP-0108-10), as previously described (https://www.protocols.io/view/fluorospot-assay-bpspmndn).

Briefly, 96-well plates (Mabtech) were coated overnight at 4°C with an antibody mixture of mouse anti-human IFN-γ (clone 1-D1K, Mabtech) and mouse anti-human IL-5 (clone TRFK5, Mabtech). 10^5^ harvested cells were plated in triplicate in coated Fluorospot plates along with either the megapools (5 μg/mL), or 10 μg/mL phytohemagglutinin (PHA) (positive control) or DMSO (negative control) and incubated for 22 hours at 37°C in 5% CO_2_. After incubation, cells were removed and membranes were washed. Anti-human IFN-γ (7-B6-1-FS-BAM, Mabtech) and anti-human IL-5 (5A10-WASP, Mabtech), in PBS with 0.1% BSA, was added and incubated for 2 hours at room temperature. Membranes were then washed again, and secondary antibodies (anti-BAM-490 and anti-WASP-640, Mabtech) in PBS with 0.1% BSA were added to the plates and incubated for 1 hour at room temperature. Lastly, membranes were washed, incubated with fluorescence enhancer (Mabtech), and air-dried prior to reading. The number of spot-forming cells (SFCs) were identified and quantified using the Mabtech IRIS system. Responses were considered positive if they met all 3 criteria: (I) DMSO background subtracted spot forming cells per 10^6^ were 100 or greater, (II) stimulation index of 2 or more compared with DMSO controls, (III) p ≤ 0.05 by Student’s t test or Poisson distribution test. Samples that did not fulfill these criteria was assigned 100 SFC/10^6^ PBMCs, the previously determined lower limit of the assay^9^. Pooled IFNγ or IL-5 values for T cell reactivity towards α-syn and PINK1 was created for donors screened for T cell reactivity towards both peptide pools. Of note, samples that not have a detectable positive response (see three criteria above) were treated as 0 during the creation of the pooled values. For samples with negative responses towards both antigens were assigned 100 SFC/10^6^ PBMCs.

### Statistical analyses

Graphs were created, and statistical analyses were performed using GraphPad Prism (GraphPad Prism, v10, RRID:SCR_002798). Two-tailed Kruskal-Wallis tests followed by an uncorrected Dunns’ test were used to compare between groups.

## Supporting information

Key Resource Table

## Data availability

Source data for all figures are provided in the Supporting data value file, and all Fluorospot data has been uploaded to Zenodo (https://doi.org/10.5281/zenodo.14934123), and this Tier 1 data is openly available from PPMI (https://www.ppmi-info.org/access-data-specimens/download-data; RRID:SCR_006431, Project ID: 245). Clinical data used in preparation of this article for participants obtained from PPMI is openly available as Tier 1 data from PPMI, and was obtained in January 2025 from the PPMI database (https://www.ppmi-info.org/access-data-specimens/download-data; RRID:SCR_006431). For up-to-date information on the study, visit http://www.ppmi-info.org.

## Code availability

No codes were used within this study.

## Acknowledgement

Supported by LJI & Kyowa Kirin, Inc. (KKNA-Kyowa Kirin North America), the Swedish Research Council (grant references 2024-00175 to E.J.), Aligning Science Across Parkinson’s (ASAP-000375 to CSLA and DS) through the Michael J. Fox Foundation for Parkinson’s Research (MJFF). For open access, the authors have applied a CC-BY public copyright license to all Author Accepted Manuscripts arising from this submission. This work was also supported by the National Institute of Neurological Disorders and Stroke of the NIH R01NS095435 (to AS and DS), the Freedom Together Foundation (to DS), and the WHAM Investigator’s fund (to CSLA). Part of the patient data and samples received from MNI were obtained from the Quebec Parkinson Network (QPN; https://rpq-qpn.ca/). The investigators of the QPN contributed to the design and implementation of the QPN and provided data, but did not participate in the execution, analysis or writing of this report. The funders had no role in study design, data collection, analysis, publication decision, or manuscript preparation. Parts of the samples were provided by PPMI (www.ppmi-info.org/access-dataspecimens/download-data). As such, the investigators within PPMI contributed to the design and implementation of PPMI and provided data and collected biospecimens, but did not participate in the analysis or writing of this report. PPMI – a public-private partnership – is funded by the Michael J. Fox Foundation for Parkinson’s Research and funding partners, including 4D Pharma, Abbvie, AcureX, Allergan, Amathus Therapeutics, Aligning Science Across Parkinson’s, AskBio, Avid Radiopharmaceuticals, BIAL, BioArctic, Biogen, Biohaven, BioLegend, BlueRock Therapeutics, Bristol-Myers Squibb, Calico Labs, Capsida Biotherapeutics, Celgene, Cerevel Therapeutics, Coave Therapeutics, DaCapo Brainscience, Denali, Edmond J. Safra Foundation, Eli Lilly, Gain Therapeutics, GE HealthCare, Genentech, GSK, Golub Capital, Handl Therapeutics, Insitro, Jazz Pharmaceuticals, Johnson & Johnson Innovative Medicine, Lundbeck, Merck, Meso Scale Discovery, Mission Therapeutics, Neurocrine Biosciences, Neuron23, Neuropore, Pfizer, Piramal, Prevail Therapeutics, Roche, Sanofi, Servier, Sun Pharma Advanced Research Company, Takeda, Teva, UCB, Vanqua Bio, Verily, Voyager Therapeutics, the Weston Family Foundation and Yumanity Therapeutics.

## Author contributions

C.S.L.A., D.S., and A.S. participated in the design and direction of the study, E.J., A.Fre., G.P.W., T.M., and J.S. performed and analyzed the experiments. A.Fra., N.K.T, I.L., J.G.G., R.N.A., D.G.S., A.W.A., N.S., E.A.F., and R.B.P. recruited participants and performed clinical evaluations. E.J., D.S., C.S.L.A., and A.S. wrote the manuscript. All authors read, edited, and approved the manuscript before submission.

