## Supplementary Information for "T cell responses towards PINK1 and α-synuclein are elevated in prodromal Parkinson’s disease"

**Figures**

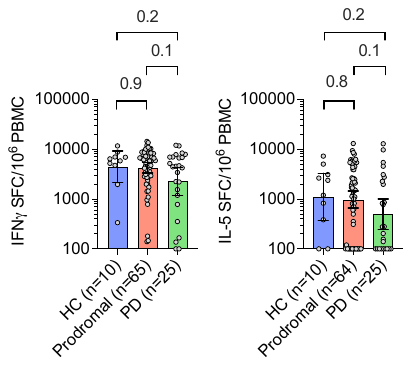

**Supplemental Figure S1. No difference in T cell responses to EBV between the three groups**. Magnitude of IFNγ (left panel) and IL-5 (right panel) mediated T cell responses towards EBV in healthy controls (HC), prodromal, and Parkinson’s disease (PD) donors. Each circle represents an individual donor. P-values from two-tailed Kruskal-Wallis tests followed by an uncorrected Dunns’ test, and geometric mean ± 95% confidence interval are shown.

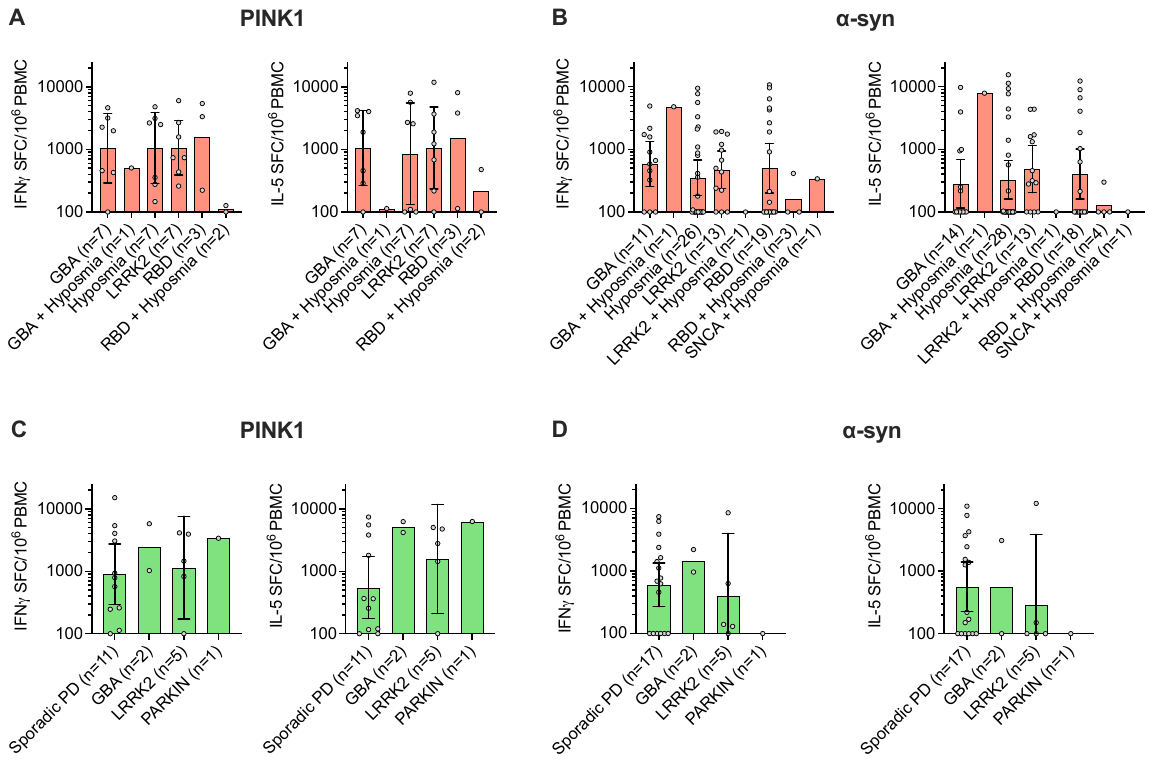

**Supplemental Figure S2. Separation of prodromal into subgroups**. Magnitude of IFNγ and IL-5 mediated T cell responses towards PINK1 (**A**) and ⍺-synuclein (⍺-syn; **B**) in prodromal donors, divided into subgroups based on Parkinson’s disease (PD)-associated genetic mutations in GBA or LRRK2 genes, diagnosis of conditions known to commonly precede PD diagnosis, hyposmia and REM sleep behavior disorder (RBD), a combination of genetic mutations and hyposmia, or a combination hyposmia and RBD. Each circle represents an individual donor. Geometric mean is shown, and for groups with five or more donors ± 95% confidence interval is shown.

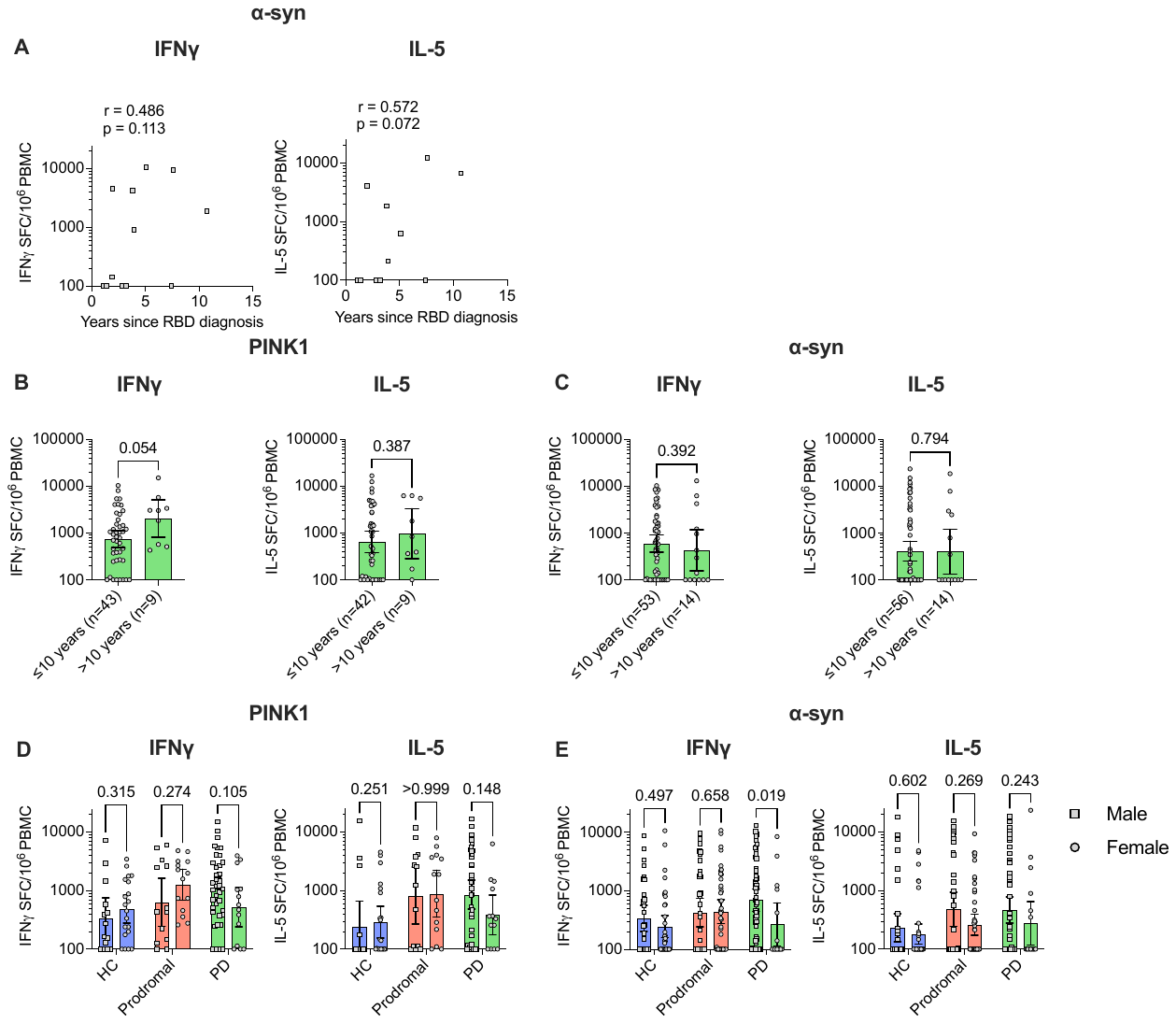

**Supplemental Figure S3. Inter-group comparison based on time since diagnosis and biological sex.** Association between IFNγ (left panel, n=12) and IL-5 (right panel, n=11) mediated T cell responses towards ⍺-synuclein (⍺-syn) and years since diagnosis in RBD donors (**A**). Comparison of the magnitude of IFNγ (left panel) and IL-5 (right panel) mediated T cell responses towards PINK1 (**B**) and ⍺-syn (**C**) in PD donors, divided into participants who donated samples within or after 10 years of diagnosis. Inter-group comparison of the magnitude of IFNγ (left panel) and IL-5 (right panel) mediated T cell responses towards PINK1 (**D**) and ⍺-syn (**E**) between male (square symbols, left bar) and female (circle symbols, right bar) participants. Correlation is indicated by Spearman r and p-value. For group-wise comparisons, p-values from two-tailed Mann-Whitney tests and geometric mean ± 95% confidence interval are shown.
